# Solution structure, dynamics and fragment binding of unbound MERS-CoV nsp10

**DOI:** 10.64898/2026.09.24.754126

**Authors:** Danni Dong, Frank Kozielski, Christopher A. Waudby

## Abstract

Middle East Respiratory Syndrome Coronavirus (MERS-CoV) poses a significant public health threat, with a fatality rate of 37% and no approved therapeutics (World Health Organization, 2026). Non-structural protein 10 (nsp10) is an essential cofactor that activates both the nsp14 3’-5’ exoribonuclease (ExoN) activity required for RNA proofreading and the nsp16 2’-O-methyltransferase (2’-O-MTase) implicated in viral RNA cap formation. Despite its functional importance, the unbound structure and dynamics of MERS-CoV nsp10 in solution remain uncharacterised. Here we report a near-complete NMR backbone and sidechain assignment and characterise the solution structure and dynamics of the protein by NMR. In contrast to the folded α1 helix observed in crystal structures of coronavirus nsp10, we find that this region is intrinsically disordered in solution, with residues 10-22 undetectable under standard conditions and further evidenced by pH titration, temperature-dependent NMR, and CLEANEX-PM experiments. Analysis of NOE contacts confirmed that the core adopts the conserved coronavirus nsp10 fold, consistent with the AlphaFold-predicted structure. ^15^N backbone Relaxation measurements indicated a rigid, well-ordered core with only localised flexibility, despite a relatively low proportion of secondary structure elements, and CPMG and CEST experiments detected no conformational exchange on the μs-ms timescale. Building on this structural and dynamic characterisation, we explored the ligandability of nsp10 by ^19^F NMR fragment screening. Screening of a 463-compound library identified 23 initial hits (4.97% hit rate), of which 20 were confirmed by ^15^N SOFAST-HMQC and eight gave quantifiable affinities by MST (K_d_ 0.5-6.9 mM), the remainder being too weak for reliable determination. Chemical shift perturbations clustered near functional surfaces of the folded core, indicating that MERS-CoV nsp10 is ligandable and providing chemical starting points for antiviral development.

## Introduction

Middle East respiratory syndrome (MERS), caused by MERS-CoV, first emerged in Saudi Arabia in April 2012 and continues to cause regional outbreaks with a particularly high case fatality rate of 37% (World Health Organization, 2026), which is substantially higher than that observed for SARS-CoV (10%) or SARS-CoV-2 (1%) (Petersen, Koopmans et al. 2020), albeit with a significantly lower infection rate. Despite this ongoing public health threat, MERS-CoV remains understudied compared to SARS-CoV-2. Currently no specific antiviral therapies or vaccines have been approved for clinical use against MERS-CoV. This situation emphasises the urgent need for targeted research into MERS-CoV molecular biology and drug discovery studies.

Nsp10 functions as an essential cofactor that activates both the nsp14 3’-5’ exoribonuclease (ExoN) activity required for proofreading and nsp16 2’-O-methyltransferase (2’-O-MTase) activity for viral RNA cap formation. The proofreading mechanism performed by the ExoN domain secures replication fidelity (Eckerle, Becker et al. 2010) and maintains the viral replication level (Ogando, Zevenhoven-Dobbe et al. 2020). The proofreading mechanism is also involved in the development of resistance to certain nucleotide analogue inhibitors such as Remdesivir, as the ExoN domain can excise incorporated nucleotide analogues from nascent viral RNA (Agostini, Andres et al. 2018). In SARS-CoV, nsp10 can stimulate the ExoN activity by more than 35-fold (Bouvet, Imbert et al. 2012). Nsp16 acts as an nsp10-dependent 2’-O-MTase to form the viral RNA cap structure, a process essential for translation efficiency and evasion of host innate immune recognition (Wang, Sun et al. 2015). Evidence in SARS-CoV has shown that nsp16 activity is completely abolished without nsp10 binding (Chen, Cai et al. 2009). These findings highlight the potential of nsp10 as a drug target, and the importance of understanding nsp10 structure and dynamics.

Previous crystallography studies have reported the MERS-CoV nsp10-nsp16 complex (PDB IDs 5YN5 and 5YN6), while the unbound nsp10 structure remains unavailable. Previous crystallisation attempts in our laboratory, screening commercially available conditions at both room temperature and 4 °C across a range of protein concentrations, did not yield crystals of the unbound protein. Solution NMR spectroscopy was therefore used, which additionally allows the dynamics of the protein to be characterised in solution.

In this study, we obtained near-complete backbone and sidechain NMR assignments of MERS-CoV nsp10 except the N-terminal region, characterised protein structure and dynamics across multiple timescales and investigated the conformational properties of the important N-terminal region essential for nsp14 and nsp16 binding. This project provides first structural insights of solution-state nsp10 with comprehensive dynamics analysis. The NMR-derived structural and dynamic characterisation reported in this study reveals key features of unbound nsp10, including N-terminal disorder and backbone rigidity in the protein core, providing insight of how this essential cofactor recognises and activates its partner proteins. These findings may also guide future structure-based approaches to disrupt functionally critical protein-protein interactions. While previous studies have explored nsp10 as a drug target in SARS-CoV-2, the ligandability of MERS-CoV nsp10 has not been directly investigated. Here, fragment screening was conducted using ligand-observed ^19^F-NMR, followed by protein-observed NMR validation, binding site mapping and affinity quantification by MST, to evaluate MERS-CoV nsp10 as a target for small molecule intervention.

## Methods

### Construct design

The protein sequence of MERS-CoV nsp10 was obtained from the NCBI reference sequence entry YP_009944301.1. The corresponding cDNA coding for amino acids 10 to 131 was codon-optimised for expression in *E. coli*, synthesised, and subcloned into the ppSUMO-2 vector (GenScript) using NcoI and XhoI restriction sites. The resulting construct encoded an N-terminal His-tag, followed by a SUMO tag, a ULP1 protease cleavage site, and the nsp10 coding sequence. After ULP1 protease cleavage, the final construct contained three non-specific N-terminal residues (TMG) followed by the MERS-CoV nsp10 sequence (residues 10-131), yielding a total construct length of 125 residues (see supplementary sequences).

### Sample preparation

The expression construct was transformed into Rosetta 2 (DE3) competent cells. For ^13^C, ^15^N-labelled protein, M9 medium containing ^15^N-ammonium chloride and U-^13^C_6_ D-glucose as the sole nitrogen and carbon sources, respectively, was used. For unlabelled protein, LB medium was used. Cultures were grown at 37°C to an OD_600_ of 0.7, induced with 0.5 mM IPTG and expressed at 20°C for 20 h. Cells were harvested by centrifugation at 6,000 × g for 20 min at 4 °C and resuspended in buffer A (50 mM sodium phosphate, pH 8.0, 300 mM NaCl, 10 mM imidazole and 1 mM PMSF), flash-frozen in liquid nitrogen, and stored at – 80°C.

Cells were lysed by sonication and clarified by centrifugation at 20,000 rpm for 60 min at 4°C. The supernatant was loaded onto a 5 mL HisTrap FF crude column (Cytiva) pre-equilibrated with Buffer A. After washing with 50 column volumes (CVs) of Buffer B (50 mM Tris-HCl pH 8.0, 300 mM NaCl, 20 mM imidazole), bound protein was eluted with Buffer C (50 mM Tris-HCl pH 8.0, 300 mM NaCl, 300 mM imidazole). The His-SUMO tag was removed by overnight dialysis at 4°C with ULP1 protease (1 mg ULP1 per 50 mg protein) in buffer E (50 mM Tris-HCl pH 8.0, 300 mM NaCl), followed by a second Ni-NTA column to remove the cleaved His-SUMO tag. Purified protein was exchanged into NMR buffer (10 mM sodium phosphate pH 8.0, 100 mM NaCl, 1 mM DTT) and concentrated to 50-200 μM. Purity was assessed by SDS-PAGE and protein integrity was confirmed by LC-MS.

For MST assays, His-tagged protein was required for fluorescent labelling; purification was therefore performed identically but without ULP1 cleavage. The protein was concentrated to 10 mg/mL, snap-frozen, and stored at –80 °C.

### NMR Spectroscopy

NMR samples were supplemented with 10% (v/v) D_2_O and 0.001% (w/v) DSS as an internal chemical shift reference. Spectra were recorded at 298 K unless otherwise stated on Bruker spectrometers at 600, 700, 800 and 950 MHz equipped with cryogenic triple-resonance probes (UCL School of Pharmacy; MRC Biomedical NMR Centre, Francis Crick Institute). All NMR data were processed with TopSpin version 4.4.1 (Bruker Biospin) and analysed with CCPNmr Analysis version 3 (Skinner, Fogh et al. 2016). Acquisition parameters are summarised in Table S1.

Sequential backbone assignments were obtained using standard triple-resonance experiments (BEST-HNCACB, HNCACO, HNCO, HNCOCACB) (Lescop, Schanda et al. 2007) and ^1^H-^15^N SOFAST-HMQC at 600 MHz. Sidechain assignments were obtained using (H)CCH-TOCSY, H(C)CH-TOCSY (Lescop, Schanda et al. 2007) and ^1^H-^13^C HSQC experiments at 800 MHz. Structural constraints were derived from ^15^N-NOESY (600 MHz, 80 ms mixing time) and ^13^C-NOESY (700 MHz, 160 ms mixing time) experiments. For backbone assignment at 283 K, BEST-HNCOCA, HNCA, HNCO, HNCOCACB were recorded at 800 MHz (Table S1).

The structure of MERS-CoV nsp10 was predicted using the AlphaFold Server (Abramson, Adler et al. 2024). SPANR analysis (Williams, Gagnon et al. 2025) was performed using the AlphaFold-predicted structure, chemical shift assignments in NEF format, the ^15^N NOESY peak list and the Predicted Aligned Error (PAE) matrix, to validate the predicted fold against experimental NMR data. The ^15^N-NOESY peak list was exported from CCPNmr Analysis v3. A modified SPANR implementation accepting NEF format was developed by Dr. Gary Thompson in collaboration with the original authors. The method calculates Contact Score (CS) and Distance Score (Petersen, Koopmans et al.) heuristics, which together with the PAE serve as inputs to a pre-trained Support Vector Machine classifier that outputs the likelihood that the predicted structure has a TM-score > 0.5 relative to the true structure (Zhang and Skolnick 2004).

Secondary structure was predicted from backbone chemical shifts (^1^HN, ^15^N, ^13^C’, ^13^Cα and ^13^Cβ) using TALOS-N version 4.21 (Shen and Bax 2013) on NMRBox (Maciejewski et al. 2017). Predictions were obtained for residues 23 to 131; residues 10 to 22 were excluded as no assignments were available for this region. Predictions for residues with incomplete assignments were treated as low confidence. Secondary chemical shifts were calculated as the difference between observed ¹³Cα and ¹³Cβ chemical shifts and the corresponding random coil values.

### Relaxation dynamics and exchange studies

Heteronuclear ^15^N relaxation measurements (R_1_, R_2_ and {^1^H}-^15^N heteronuclearNOE) were performed at 298 K on both 800 MHz and 600 MHz spectrometers for model-free analysis using sensitivity-enhanced HSQC-based pulse sequences (Farrow, Muhandiram et al. 1994) on a 200 μM ^15^N-labelled sample. The R_1_ experiment incorporated temperature compensation to minimise duty-cycle heating (Yip and Zuiderweg 2005). R_1_ was measured using the relaxation delay times of 20, 100, 200, 350, 500, 750, 1000 and 1500 ms; R_2_ with delays of 16.96, 33.92, 50.88, 67.84, 84.80, 101.76, 135.68 and 169.6 ms was recorded. {^1^H}-^15^N heteronuclearNOE was determined from the ratio of peak intensities(I_sat_/I_0_) obtained with and without 3 s amide proton saturation.

Model-free analysis used R_1_, R_2_ and heteronuclear NOE data at 800 MHz, together with R_1_ and R_2_ at 600 MHz. The heteronuclear NOE data at 600 MHz were excluded due to systematic disagreement with other relaxation parameters, likely arising from temperature fluctuations during the extended acquisition period.

^15^N CPMG relaxation dispersion measurements were conducted at 700 MHz (Hansen, Vallurupalli et al. 2008) with T_relax_ of 40 ms and v_CPMG_ frequences of 25, 50, 75, 100, 125, 150, 175, 200, 225, 250, 300, 350, 400, 450, 500, 600, 700, 800, 900, 1000 Hz (20 points). ^15^N CEST (Chemical Exchange Saturation Transfer) experiments were performed at 950 MHz (Vallurupalli, Bouvignies et al. 2012) with a saturation time of 300 ms and three saturation field strengths (15, 30, and 60 Hz), sampling offests from –1200 to +1200 Hz in 50 Hz increments.

All relaxation and exchange data were analysed using NMRAnalysis.jl (https://github.com/waudbylab/NMRAnalysis.jl). Modelfree analysis were conducted with FASTModefree (Cole and Loria 2003) on NMRBox (Maciejewski, Schuyler et al. 2017).

### Temperature, pH dependence, and amide exchange studies

Temperature-dependent ^1^H-^15^N SOFAST-HMQC spectra were acquired from 273 K to 298 K in 5 K increments on a 150 μM protein sample at 800 MHz. For pH titration, the same sample was supplemented with 100 μM HEPES as an internal pH indicator, and ^1^H-^15^N SOFAST-HMQC spectra were recorded at pH values of 7.5, 7.0, 6.5, and 6.0 at 298 K. pH adjustments were made by titrating with small aliquots of 1 M HCl, with sample pH verified from the ^1^H chemical shift of the HEPES piperazine CH_2_ resonance using an in-house calibration curve.

CLEANEX-PM experiments (Hwang, Van Zijl et al. 1998) were performed at 800 MHz and 278 K to identify amide protons undergoing rapid solvent exchange with mixing times of 40 ms and 80 ms.

### Fragment screening

A fragment library of 463 compounds from the BioNET ^19^F library, adhering to “Lipinski rule of three”, was screened against unlabelled nsp10 (0.5 mg/ml) in 50 mM sodium phosphate pH 7.4, 100 mM sodium chloride, 50 μM DSS, 10% D_2_O. Fragment cocktails (∼30 fragments each, 50-100 μM individual concentration) were prepared alongside protein-free references using a Bruker SamplePro Tube liquid handling robot.

One-dimensional ^1^H decoupled spectra and ^19^F T_2_ relaxation measurements were recorded at 600 MHz and 298 K using a perfect-echo broadband pulse sequence with relaxation delays from 5 to 500 ms. Data were processed using NMRScreen.jl (https://github.com/waudbylab/NMRScreen.jl). Fragments with ΔR_2_ > 10 s^-^¹ were identified as hits.

### Fragment validation by protein-observed NMR

^15^N-labelled nsp10 (10 μM) in NMR buffer (10 mM sodium phosphate pH 8.0, 100 mM NaCl, 50 μM DSS, 10% D_2_O, 1 mM DTT) was prepared and ^1^H-^15^N SOFAST-HMQC experiments were recorded at 700 MHz and 298 K for the protein alone and in the presence of 1 mM of each fragment.

To assess the chemical shift perturbation (CSP) induced by fragments, peaks were assigned to their nearest neighbours, and Euclidean distances were calculated based on ^1^H and ^15^N chemical shift changes using the equation:

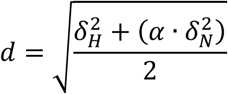

where α = 0.122, determined from the ratio of observed ^1^H and ^15^N chemical shift ranges (Williamson 2013). Significant CSPs were defined as those exceeding the mean plus one standard deviation.

### Microscale thermophoresis (MST) binding affinity measurements

Binding affinities were estimated by MST using a Monolith NT.115 instrument. Fragment stocks were prepared at 10 mM in PBST buffer (pH 7.4) and serially diluted (2-fold, 10 mM to 30 μM). Equal volumes of 20 nM His-tagged nsp10 were mixed with each concentration and incubated on ice for 1 h. Measurements were performed in triplicate and analysed using MO.Control and MO.Affinity Analysis software.

## Results

### Solution NMR reveals a structured core but an undetectable N-terminal region

The N-terminal α1 helix is consistently observed in crystal structures of unbound nsp10 and of nsp10 in complex with nsp14 from SARS-CoV and SARS-CoV-2, where it forms part of the nsp14 binding interface (Figure 1A) (Lin, Chen et al. 2021). By contrast, in nsp10-nsp16 complexes, it lies outside the binding interface (Figure 1B) (Wei, Yang et al. 2018). Consistent with the nsp14-bound and unbound structures, AlphaFold predicts a folded helix for residues N10-F19 of MERS-CoV nsp10, followed by a short loop formed by T20-P22 connecting α2 helix. Confidence in this region is nevertheless lower than for the folded core, both in the local prediction (mean pLDDT 84.6 for residues 10-22, compared with 93.6 for residues 23-131) and in its position relative to the core (averaged predicted aligned error 4.9 Å versus 2.6 Å) (Figure 1C). We therefore paid particular attention to this region in our subsequent analysis of the solution-state NMR data.

**Figure 1.**
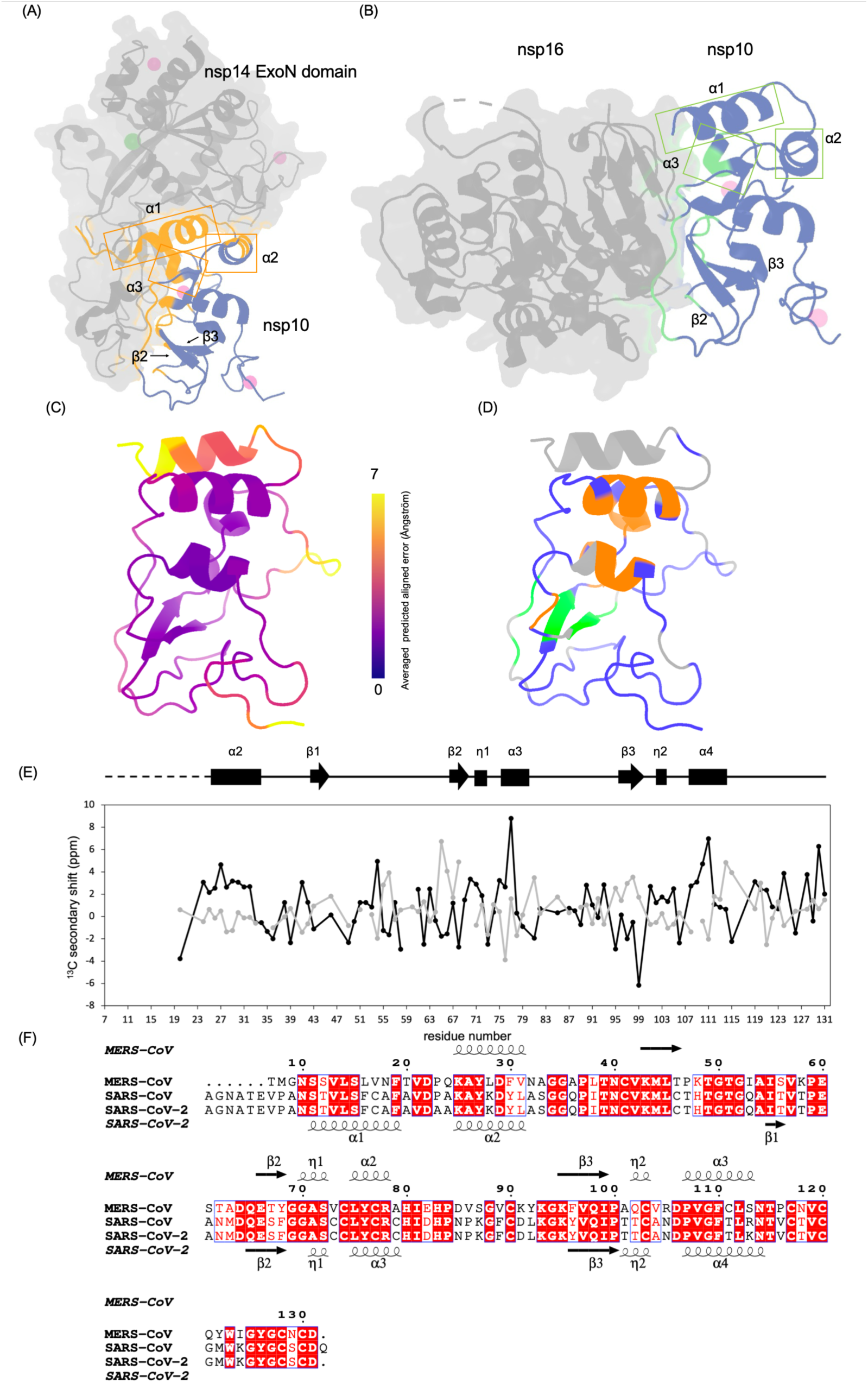
Structural context and NMR-based characterisation of MERS-CoV nsp10. (A) Crystal structure of the SARS-CoV-2 nsp14-nsp10 complex (PDB ID 7DIY), showing nsp10 (blue) bound to the nsp14 ExoN domain (grey surface). Regions of nsp10 at the nsp14 interface are highlighted in orange, including the N-terminal α1 helix. (B) Crystal structure of the MERS-CoV nsp16-nsp10 complex (PDB ID 5YN5), showing nsp10 (blue) bound to nsp16 (grey surface). Regions of nsp10 at the nsp16 interface are highlighted in green. The α1 helix lies outside the nsp16 interface. (C) AlphaFold-predicted structure of MERS-CoV nsp10 coloured by averaged predicted aligned error (PAE), from 0 Å (dark purple) to 7 Å (yellow). Purple indicates regions of higher PAE and correspondingly lower confidence in their position relative to the folded core. (D) The same AlphaFold model, in an identical orientation, coloured by NMR assignment status: assigned residues are coloured by TALOS-N secondary structure (α-helix, orange; β-sheet, green; loop, blue), and residues not detected under standard conditions (600MHz, 298 K, pH 8.0) are shown in grey. (E) Secondary chemical shift analysis. Upper: schematic representation of TALOS-N secondary structure prediction based on NMR chemical shifts. Lower: secondary chemical shift values for ^13^Cα (black) and ^13^Cβ (grey) relative to random coil values. The absence of data for residues 10-22 reflects the lack of observable signals for this region. (F) Sequence and secondary structure alignment of MERS-CoV, SARS-CoV, and SARS-CoV-2 nsp10. Identical residues are shown in white text on a red background, similar residues in red text, and different residues in black. The alignment was generated using CLUSTAL OMEGA (Sievers, Wilm et al. 2011), and secondary structure elements were added using ESPript 3.0 (Gouet, Courcelle et al. 1999). Secondary structures for SARS-CoV and SARS-CoV-2 are identical and derived from crystal structures (PDB IDs 2FYG and 6ZCT), while MERS-CoV nsp10 secondary structure represents TALOS-N predictions. Residues are numbered according to the native nsp10 sequence.

Near-complete backbone (Sattler, Schleucher et al. 1999) and sidechain (Bax, Clore et al. 1990) assignments were obtained using triple-resonance NMR experiments (Figure S1,2). No signals were detected for residues 10-22 under standard conditions (298 K, pH 8.0). Excluding this region, backbone assignment achieved 92% completion for both ^1^H and ^15^N amide signals and for ^13^C signals (91% C’, 91% C_α_, 94% C_β_), and sidechain assignments reached 85% completion for proton and 91% for carbon signals. LC-MS analysis confirmed the molecular weight of the NMR sample (13,350.22 Da) matched the theoretical value (13,350.29 Da) computed by ExPASy (Gasteiger, Gattiker et al. 2003), ruling out protein truncation or degradation as the cause of signal absence.

Chemical shift analysis with TALOS-N (Shen and Bax 2013) revealed a conserved secondary structure organisation comprising three α-helices (K25-N31, L75-R78, and P107-L112), two 3_10_ helices (G70-S72 and Q102-C103) and β-strands at E66-S68 and K95-I99 (Figure 1E). K43-T46 is predicted as β1 by TALOS-N mainly from sequence as only K43 has sufficient assignments, therefore should be interpreted with caution. Following the coronavirus nsp10 crystal structures, these helices correspond to α2, α3 and α4, while no signals were observed for the region corresponding to α1. This arrangement corresponds to SARS-CoV and SARS-CoV-2 nsp10 folding of the core region despite a sequence identity of only 59.4% (Figure 1F), indicating that the coronavirus nsp10 structural core fold is conserved in MERS-CoV.

SPANR analysis (Williams, Gagnon et al. 2025) validated the AlphaFold-predicted structure against the experimental NMR data (^15^N NOESY), yielding the following heuristic scores: Contact Score (CS) of 0.020 and Distance Score of 0.476, and a likelihood of 92.1% that the predicted structure is consistent with the NMR constraints (defined as TM-score >0.5 relative to the true structure). This validation confirms the reliability of using the AlphaFold-predicted model for interpreting the NMR relaxation and dynamics data for the well-ordered core region (residues ∼23-131). The predicted structure is consistent with MERS-CoV nsp10 crystal structural in complex with nsp16 (PDB: 5YN5). The structural conservation despite moderate sequence identity is consistent with the functional importance of nsp10, as precise positioning of interface residues is required for activation of both nsp14 and nsp16 across different coronaviruses.

### The N-terminal region exhibits intrinsic disorder in solution

In contrast to the conserved core, the N-terminal region (predicted α1 helix) behaves differently in solution. In the AlphaFold model of MERS-CoV nsp10 (Figure 1C) and in the crystal structure of the MERS-CoV nsp10-nsp16 complex (Figure 1B) (Wei, Yang et al. 2018), α1 packs against the α2 helix. No signals were detected for residues 10-22 under standard conditions (pH 8.0, 298 K), and sidechain resonances could not be assigned for this region, with the exception of a tentatively assigned T20 identified from its characteristic chemical shifts. Therefore, the expected inter-helical NOEs could not be located in the ^13^C-NOESY spectra, and the conformational state of this region was instead examined by pH titration, temperature-dependent NMR and CLEANEX-PM experiments.

pH titration from pH 7.5 to 6.0 revealed the appearance of new amide signals at lower pH, indicating reduced amide proton exchange with bulk water that made signals from previously invisible flexible regions detectable (Figure 2A). Several residues showed significant pH-dependent chemical shift changes, with the largest perturbations observed for C77, R78, A79 and H80 (Figure 2B). These residues are located at the nsp14 and nsp16 binding interface, with H80 involved in zinc coordination. However, the large perturbation of H80 is also consistent with direct protonation of the histidine imidazole side chain (pKa ∼6.0-6.5), which falls within the titration range used here; distinguishing intrinsic histidine titration from functionally relevant pH-dependent modulation of the binding interface would require further investigation.

**Figure 2.**
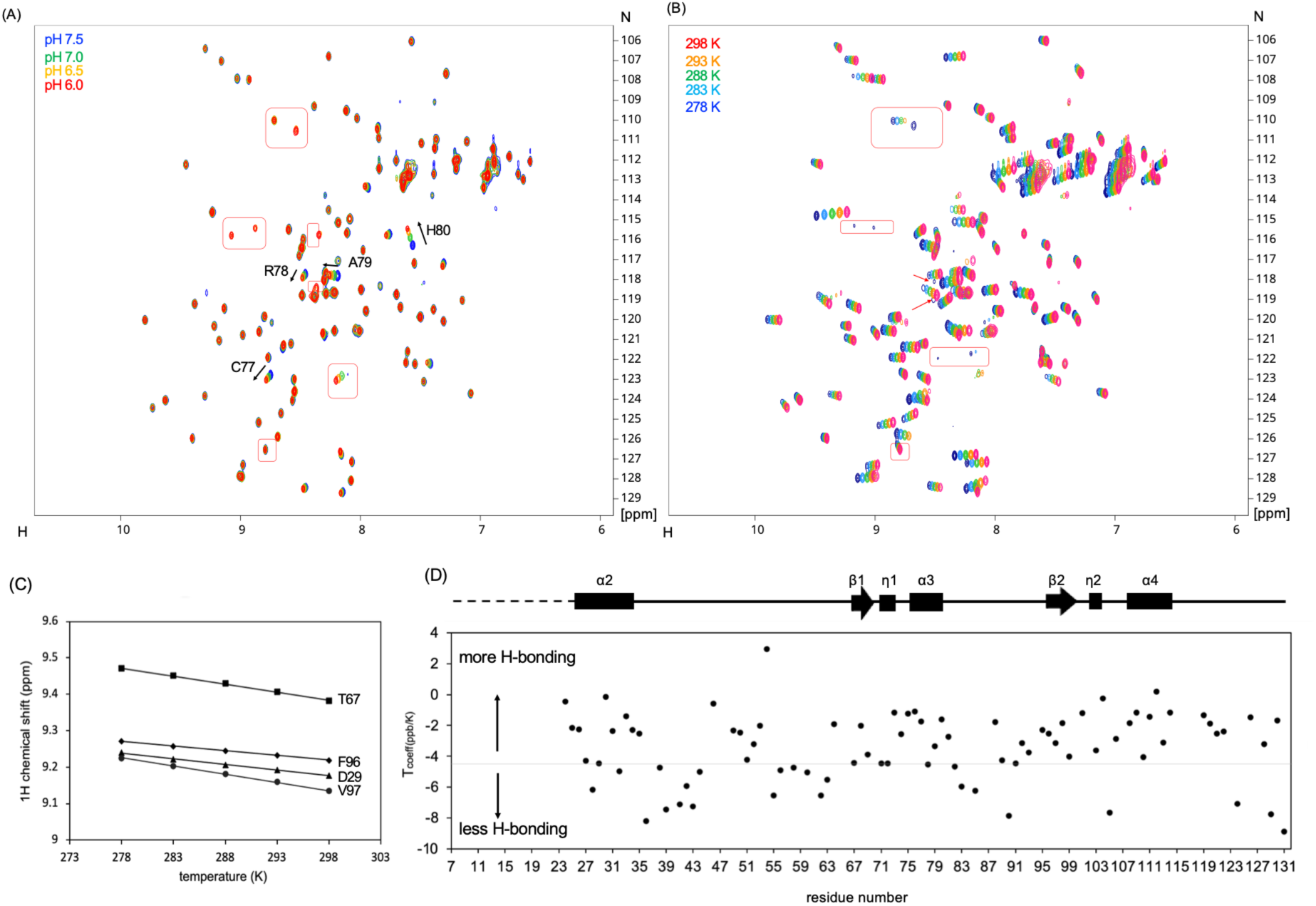
pH titration and temperature-dependent NMR analysis of MERS-CoV nsp10. (A) Overlaid ^1^H-^15^N SOFAST-HMQC spectra acquired during pH titration from pH 7.5 to 6.0 at 800 MHz, 298 K. Spectra are colour-coded by pH: blue (pH 7.5), green (pH 7.0), orange (pH 6.5), red (pH 6.0). Red boxes indicate new signals appearing at lower pH. (B) Overlaid ^1^H-^15^N SOFAST-HMQC spectra acquired from 278 K to 298 K at 800 MHz, pH 7.5. Spectra are colour-coded by temperature: magenta (298 K), orange (293 K), green (288 K), sky blue (283 K), navy (278 K). Red boxes and arrows indicate signals that appear upon temperature reduction. (C) ^1^H chemical shifts of representative residues (D29, T67, F96 and V97) plotted against temperature. Amide proton temperature coefficients (T_coeff_) were obtained from the slope of a linear fit to these data. (D) Amide proton temperature coefficients (T_coeff_ = Δδ_H_/ΔT) plotted versus residue number, with TALOS-N secondary structure elements indicated above. The dashed line marks the −4.5 ppb/K threshold; values above this line indicate hydrogen bond involvement, while values below indicate reduced hydrogen bonding and greater solvent exposure.

Temperature-dependent measurements from 278 K to 298 K provided further evidence of N-terminal disorder. As the temperature decreased, new peaks appeared in the SOFAST-HMQC spectra (Figure 2B), overlapping in position with those observed during pH titration. The appearance of these signals at reduced temperature and pH indicates that the corresponding residues undergo rapid amide proton exchange with water, consistent with their location in a highly solvent-exposed region. Amide proton temperature coefficients (T_coeff_) showed values consistent with hydrogen bond formation (> –4.5 ppb/K) for secondary structure elements (Baxter and Williamson 1997), whereas loop regions (for example, A36-L45 and I55-D63) showed more negative T_coeff_ values, indicating reduced hydrogen bonding and greater solvent exposure (Figure 2D).

CLEANEX-PM experiments at 278 K directly confirmed rapid amide proton exchange with water for the newly appearing signals. Overlay of ^1^H-^15^N SOFAST-HMQC and CLEANEX-PM spectra showed that cross peaks emerging at reduced temperature overlapped with signals in the CLEANEX-PM spectrum (Figure 3A), indicating that rapid solvent exchange accounts for signal absence under standard conditions. Some CLEANEX-PM signals matched residues with existing assignments at 298 K, including I81, G88, C90, R105, and I124, indicating that these residues also undergo considerable solvent exchange despite being observable under standard conditions.

**Figure 3.**
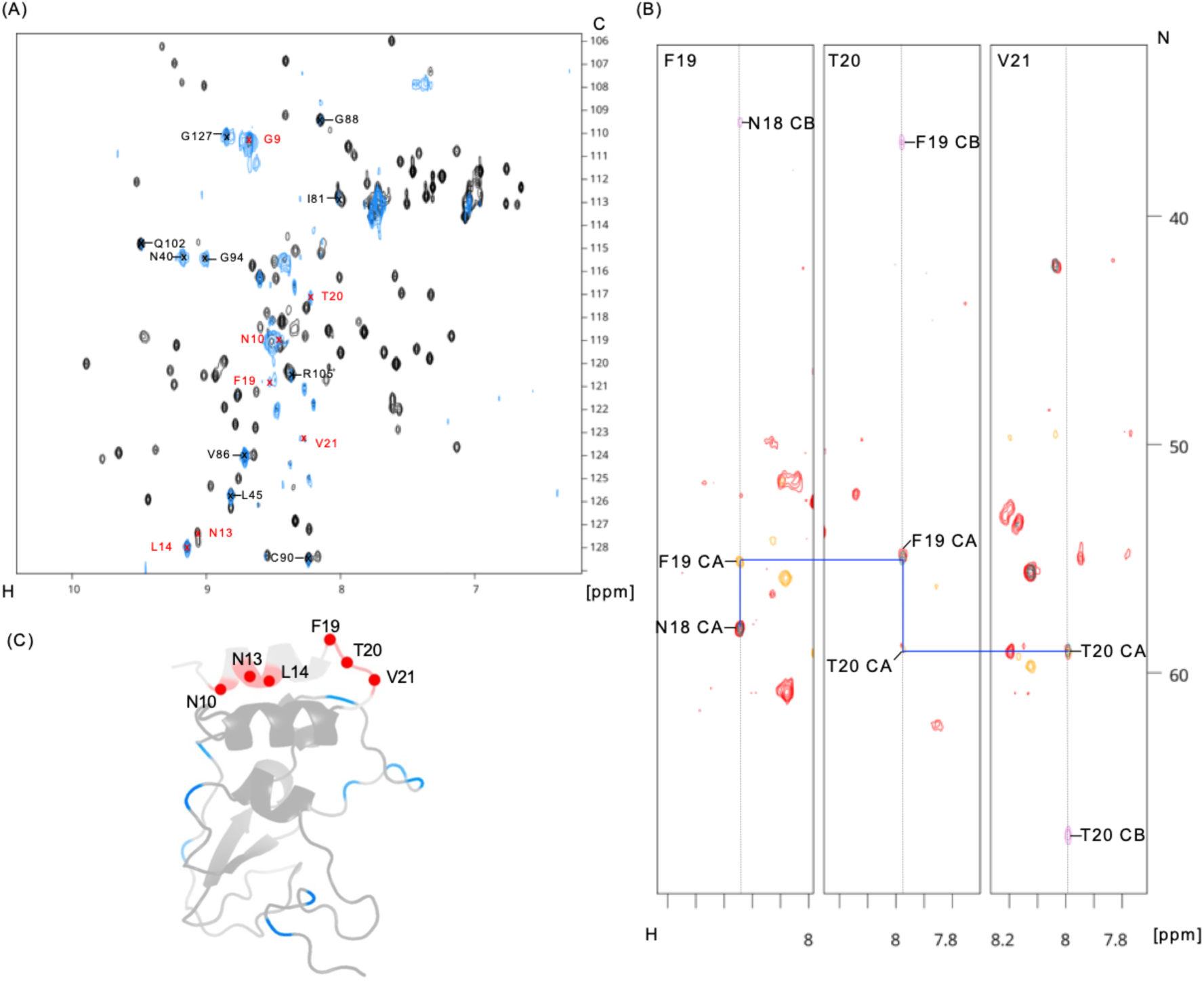
CLEANEX-PM analysis of amide proton exchange in MERS-CoV nsp10 and backbone assignment at 283 K. (A) Overlaid CLEANEX-PM spectrum (80 ms mixing time, blue) and reference SOFAST-HMQC spectrum (black) recorded at 278 K, 800 MHz. Assignments obtained at 283 K are indicated, with N-terminal residues labelled in red and loop residues labelled in black. (B) Representative strips from sequential backbone assignment of the N-terminal region at 283 K, 800 MHz: HNCA (yellow), HNCOCACB (green), and HNCOCA (red). Strips correspond to F19, T20, and V21 amide positions. (C) Residues giving CLEANEX-PM signals mapped onto the AlphaFold-predicted structure of MERS-CoV nsp10. N-terminal residues (N10, N13, L14, F19, V21) are shown in red. Other solvent-exposed residues are shown in blue. The N-terminal region (residues 10 to 22) is rendered as a transparent grey cartoon to distinguish the AlphaFold-predicted conformation from the experimentally observed residues.

To assign the unidentified signals, three-dimensional NMR experiments (HNCOCA, HNCO, HNCA, and HNCOCACB) were recorded at 283 K. Most newly assigned residues showed corresponding CLEANEX-PM signals, consistent with high solvent accessibility. These residues comprised N-terminal residues (N10, N13, L14, F19, V21) and solvent-exposed loop residues (N40, L45, V86 and G127). Some of these assignments are tentative due to the limited number of observable sequential connectivity at 283 K. This provides direct experimental evidence that the N-terminal region is disordered in unbound nsp10 in solution, in contrast to the folded helix α1 observed in nsp10-nsp14 complexes and in unbound nsp10 structures of related coronaviruses.

### Rigid backbone dynamics with localised flexibility

Despite the low content of stable secondary structure elements, _15_N relaxation experiments (R1, R_2_ and heteronuclear NOE) recorded at 298 K revealed generally rigid backbone dynamics throughout the folded region (Figure 4). Secondary structure elements exhibited heteronuclear NOE values of 0.8-0.9, characteristic of well-folded proteins.

**Figure 4.**
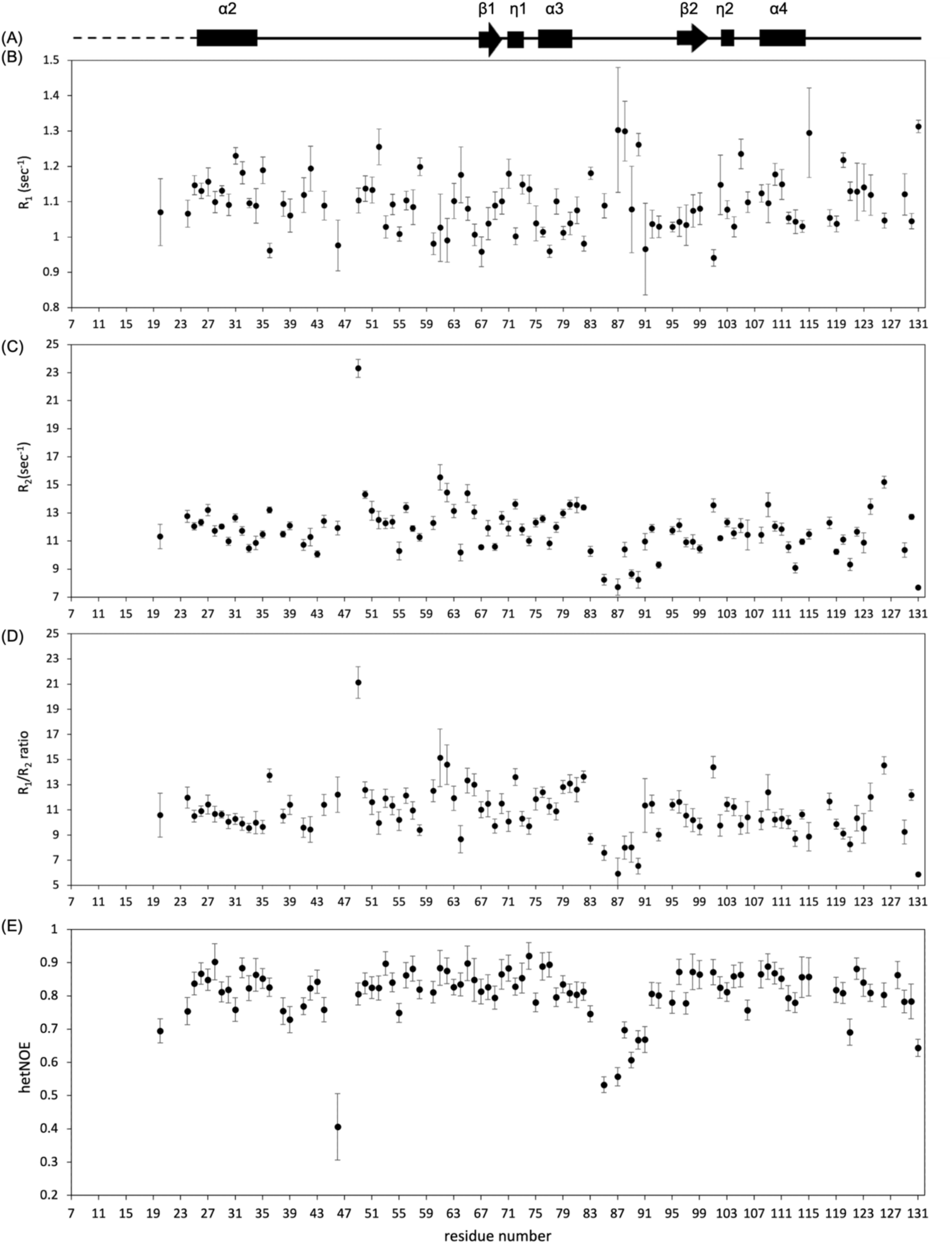
^15^N relaxation measurements of nsp10 at 800 MHz. (A) Schematic representation of TALOS-N secondary structure prediction. (B) ^15^N R_1_ values plotted versus residue number. (C) ^15^N R_2_ values plotted versus residue number. (D) R_2_/R_1_ ratios plotted versus residue number. (E) {^1^H}-^15^N heteronuclear NOE values plotted versus residue number.

Model-free analysis (Lipari and Szabo 1982) was performed to separate local flexibility from global tumbling dynamics. An axially symmetric diffusion tensor was optimised, giving τm = 7.811 ns and D_ratio_ = 1.215. Of the 90 residues with assignments, 58 were fitted to model-free models: 45 to model 1 (S^2^ only), 8 to model 2 (S^2^ and τₑ), and 5 to model 3 (S^2^ and R_ex_). Many loop residues could not be fitted to any model, likely reflecting inaccuracies in NH bond vector orientations within the AlphaFold-predicted structure in these less well-defined regions. Order parameters (S^2^) were generally greater than 0.8 for secondary structure elements, indicating a predominantly rigid backbone characteristic of a well-folded protein and consistent with the conserved globular fold observed across coronavirus nsp10 homologues (Figure 5A).

**Figure 5.**
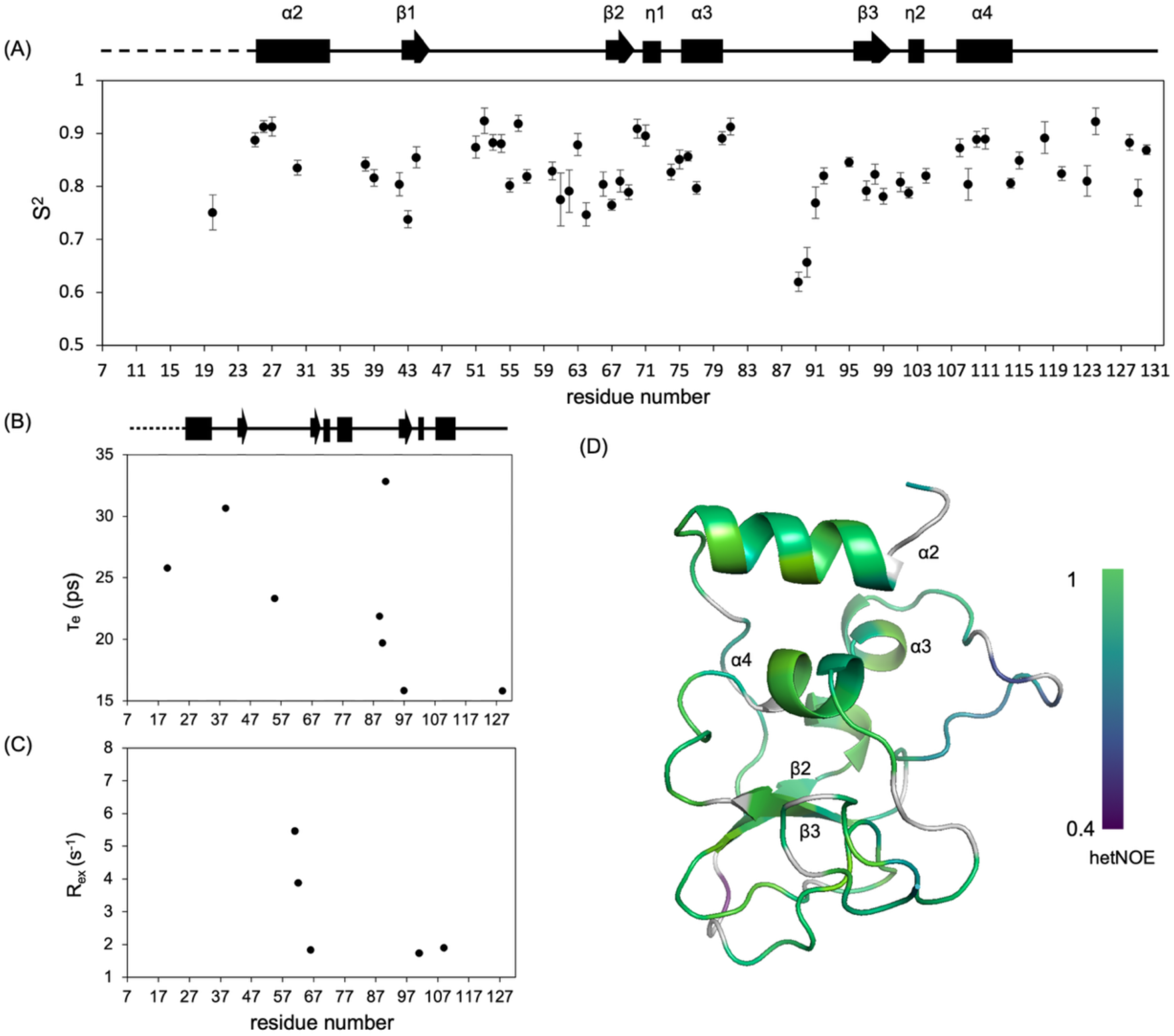
Model-free dynamic analysis for MERS-CoV nsp10. (A) Order parameter S^2^ plotted versus residue number. (B) Effective local correlation time τ_e_ plotted versus residue number. (C) Conformational exchange contribution R_ex_ plotted versus residue number. Secondary structure elements derived from TALOS-N are indicated at the top of each panel. (D) Structural model of MERS nsp10 colour-coded by amplitude of internal motion, with increasing green intensity indicating greater rigidity.

Although model-free analysis identified non-zero Rₑₓ contributions for several residues, these do not appear to reflect genuine conformational exchange. The small Rₑₓ values (< 1-2 s⁻¹) observed for E66, A101 and G109 lie within the range attributable to residue-specific variations in ^15^N chemical shift anisotropy or to limitations in model selection (Fushman 2012), while S61 and T62 displayed larger values (4-6 s⁻¹). CEST and CPMG experiments, however, detected no exchange for any of these residues. These Rₑₓ contributions therefore likely represent analytical artefacts or dynamics on timescales outside the sensitivity window of relaxation dispersion methods.

Despite this overall rigidity, localised flexibility was observed in specific regions. The loop connecting helix α3 and strand β3 (residues H83-C90) displayed notably reduced R_2_ values, order parameters and heteronuclear NOE values, indicating enhanced mobility on ps-ns timescales (Figure 5D). CLEANEX-PM experiments independently confirmed this flexibility, with residues in this region showing rapid solvent exchange. This loop lies adjacent to the nsp14-binding interface and contains the zinc-coordinating residues H83 and C90. Residues within the β-sheet of the lower subdomain (residues 66-68 and 950-99) also displayed lower order parameters than the α-helices of the upper subdomain (residues 25-31, 75-78 and 108-113), suggesting greater backbone flexibility in the lower subdomain. The functional significance of this flexibility remains to be established, but it may facilitate conformational adjustments required for optimal positioning of interface residues during partner binding.

The C-terminal region displayed unexpected rigidity compared with typical protein termini, which usually exhibit increased mobility owing to reduced structural constraints. Temperature coefficient analysis showed that C-terminal loop residues exhibited relatively high (less negative) T_coeff_ values (Figure 2D), consistent with stable hydrogen bonding networks that constrain conformational freedom in this region.

To probe conformational exchange on the ms timescale, CEST and CPMG relaxation dispersion experiments were performed for all assigned residues. Residues on helix α2 were of particular interest, as in crystal structures of nsp10-nsp14 complexes and unbound nsp10 the N-terminal α1 helix makes direct contacts with α2, we speculated that transient folding or unfolding of an α1 helix would change the chemical environment of neighbouring α2 residues and produce measurable exchange contributions. However, CEST and CPMG profiles for adjacent α2 residues, such as A33 and F30, showed no evidence of exchange (Figure 6). This indicates the absence of α1 signals does not arise from signal broadening of a folded but dynamic helix in the intermediate regime. Indeed, no significant exchange was detected for any residue across the folded region, indicating that nsp10 does not undergo detectable conformational exchange within the timescale window probed (approximately 0.3-10 ms for CPMG and ∼3-100 ms for CEST). Faster motions beyond these detection windows, or exchange involving sparsely populated states, may still exist.

**Figure 6.**
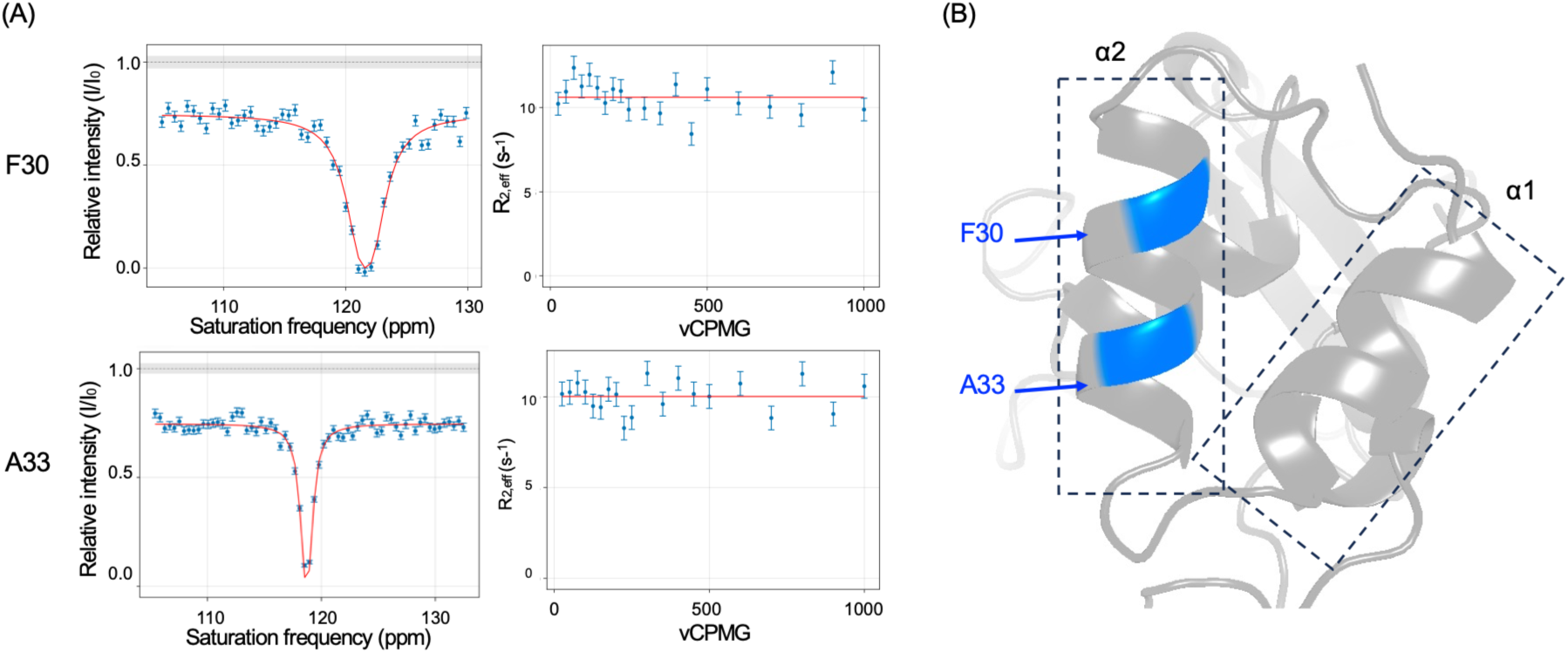
Representative CEST and CPMG profiles for α2 helix residues at 700 MHz, 298 K. (A) CEST profiles (left, 60 Hz saturation field, 300 ms saturation time) and CPMG relaxation dispersion profiles for residues F30 and A33. Red lines indicate fits to a null model (no exchange). (B) Locations of F30 and A33 (blue) mapped onto the AlphaFold-predicted structure of MERS-CoV nsp10, highlighting their positions on helix α2 adjacent to the predicted N-terminal α1 helix.

### Implications for nsp10-partner recognition

Our finding contrasts with crystal structures of coronavirus nsp10, in which the N-terminal region consistently forms the α1 helix in unbound nsp10 and in nsp10-nsp14 complexes of SARS-CoV and SARS-CoV-2. Notably, solution NMR characterisation of unbound SARS-CoV-2 nsp10 also reports α1 as helical (Kubatova, Qureshi et al. 2021), under conditions comparable to those used here (pH 7.5, 298 K). The disorder we observe for MERS-CoV nsp10 in solution therefore reflects a genuine difference between the two proteins. In nsp10-nsp16 complexes, α1 additionally adopts variable conformations (retained in 5YN5, reoriented in 6WVN, or undetectable in 7C2I), indicating that α1 is not consistently stable and may be influenced by crystal packing.

The structural basis for this species difference is not fully resolved by the present analysis. Several non-conservative substitutions at the α1-core interface, including loss of the α1-core cysteine pair (C17/C79) and of the Y30 aromatic contact, may alter the packing between α1 and the folded core. However, many substitutions within α1 itself are conservative, so the determinants of α1 disorder in MERS-CoV nsp10 remain to be established and would require experimental investigation, for example by reciprocal mutagenesis between MERS-CoV and SARS-CoV-2.

Previous functional studies revealed that the N-terminal helix α1 displays partner-specific requirements. Deletion of α1 completely abolishes binding to nsp14 and nsp14-ExoN, demonstrating that this region is essential for complex formation with nsp14 (Lin, Chen et al. 2021). However, the same deletion retains full binding capacity for nsp16, indicating that α1 is dispensable for interaction with nsp16. This functional distinction is consistent with the observation that α1 is in direct contact with nsp14 but is located on the opposite side of nsp10 from the nsp16 interface (Figure 1A, B). No structure of the MERS-CoV nsp10-nsp14 complex is currently available. In the SARS-CoV and SARS-CoV-2 complexes, α1 lies directly within the nsp14 interface, and nsp14 adopts a different conformation of its lid subdomain when bound to nsp10, which is associated with release of exonuclease activity (Czarna, Plewka et al. 2022). This functional requirement constrains the binding mode, indicating that the helical conformation of α1 is itself required for nsp14 binding and activation. CPMG relaxation dispersion and CEST experiments detected no significant conformational exchange on the ms timescale, including for α2 residues such as A26 that lie adjacent to α1. Combined with the intrinsic disorder observed for the N-terminal region, this indicates that unbound nsp10 does not populate a folded α1 conformation to a level detectable within the ms window, consistent with an induced-fit process in which α1 folds upon partner engagement. However, conformational selection mechanism cannot be excluded, since a pre-existing folded state present at low population, or exchanging outside the timescale probed here, would not be detected in these experiments.

### Fragment-based screening establishes nsp10 ligandability

Fragment screening was conducted using a library of 463 fluorine-containing compounds. Binding was assessed by monitoring changes in ^19^F transverse relaxation rates (ΔR_2_) upon protein addition (Figure 7, S3), as fragment binding to the macromolecule significantly increases rotational correlation time. Using a ΔR_2_ threshold of 10 s^-1^, 23 fragments were identified as hits, giving a hit rate of 5.0 %. Of these, 20 were confirmed by at least one orthogonal method (protein-observed NMR and/or MST), giving a validated hit rate of 4.32%.

**Figure 7.**
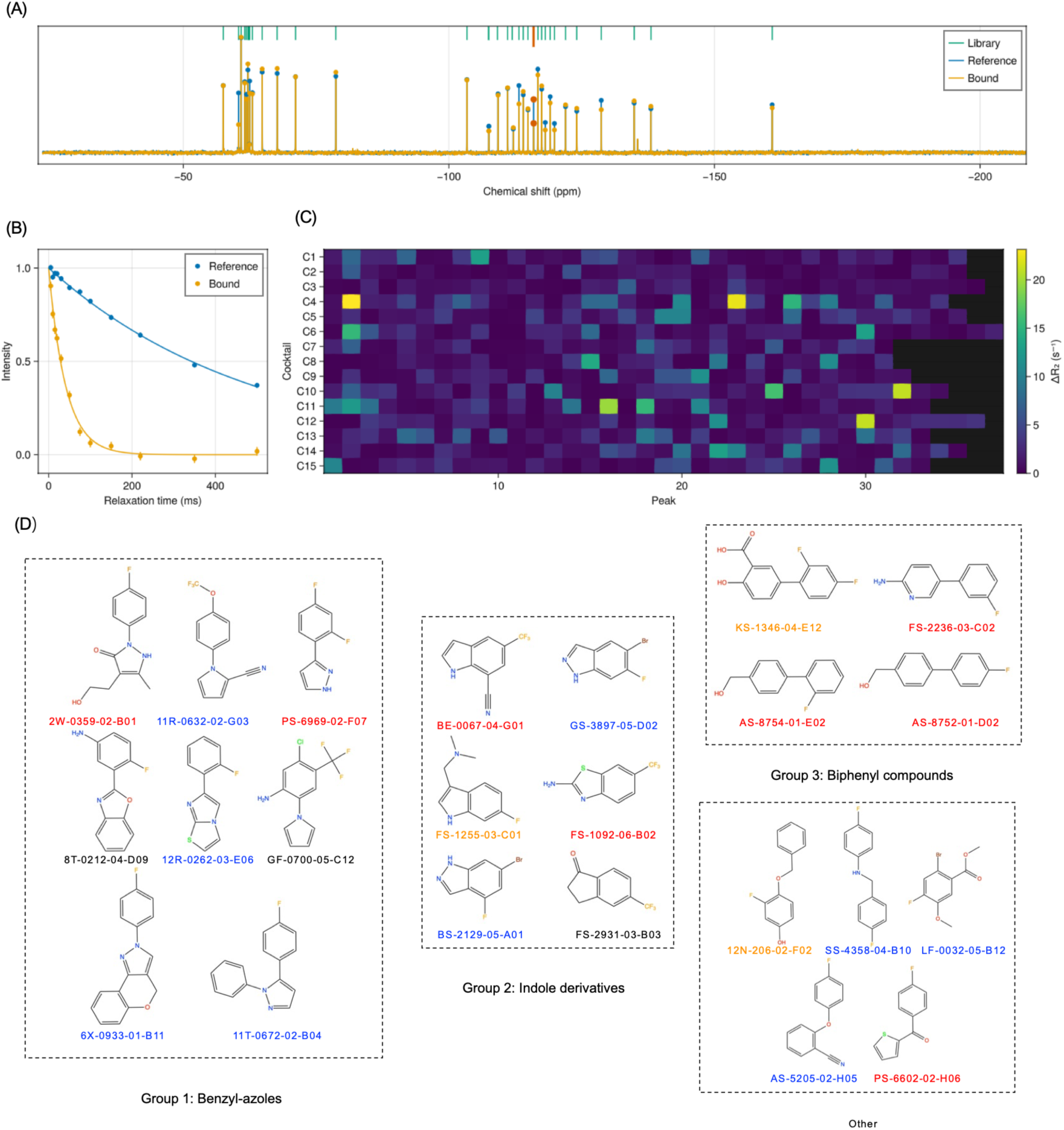
^19^F-NMR fragment screening of MERS-CoV nsp10 (600 MHz, 298 K). (A) Representative ^19^F-NMR spectrum of a fragment cocktail (C4), showing the library reference (Abramson, Adler et al.), the reference spectrum without protein (blue) and the spectrum in the presence of nsp10 (orange). Fragment binding increases the ^19^F transverse relaxation rate, reducing peak intensity. (B) ^19^F transverse relaxation decay for a representative binding fragment (BS-2129-05-A01), showing faster decay in the presence of nsp10 (orange) than in its absence (blue), corresponding to ΔR_2_ = 22.4 s^-1^. (C) Heatmap of ΔR_2_ values across all fragment cocktails (C1-C15) and ^19^F peaks. (D) Structures of the identified fragment hits, grouped by scaffold into benzyl-azoles (Group 1), indole derivatives (Group 2) and biphenyl compounds (Group 3), with remaining hits shown separately. Fragments are colour-coded by validation status: red, validated by both protein-observed NMR and MST; orange, chemical shift perturbations observed but K_d_ beyond the MST quantification range (> 10 mM); blue, perturbations observed but MST data could not be fitted; black, not validated by either method.

Protein-observed NMR was used to validate binding and to map the perturbed residues. SOFAST-HMQC spectra of ^15^N-labelled nsp10 were acquired for each hit, and three patterns of behaviour were observed: chemical shift perturbations characteristic of fast exchange, signal broadening or disappearance indicating intermediate exchange, and signal intensity enhancement. For example, fragment FS-1092-06-B02 induced chemical shift perturbations in residues T39, L75, C77, K91, Y92 and K93, which cluster spatially on the protein surface (Figure 8A).

**Figure 8.**
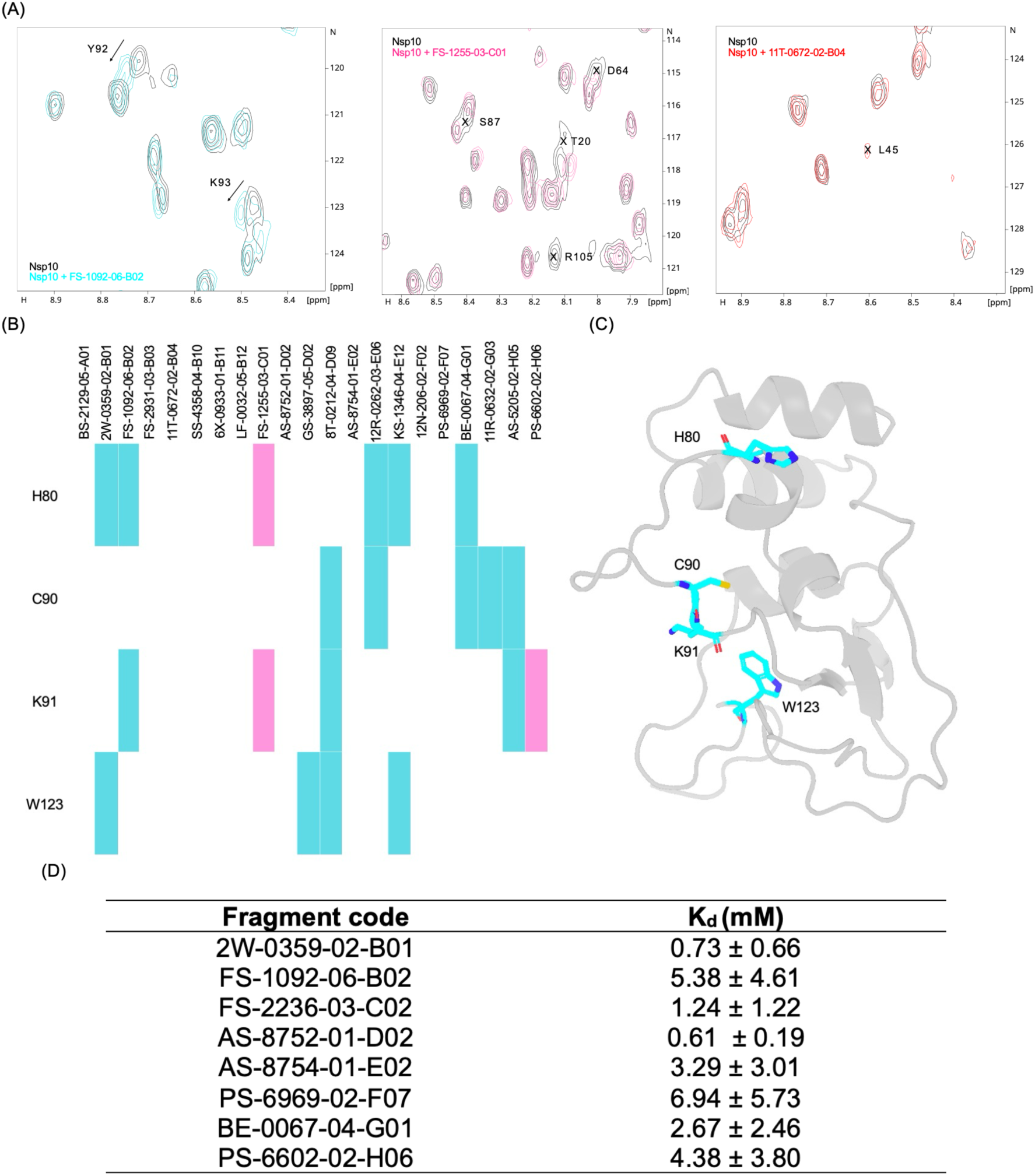
Protein-observed NMR validation, binding-site mapping and affinity determination. (A) selected regions of overlaid ^1^H-^15^N SOFAST-HMQC spectra (700 MHz, 298 K) of nsp10 in the absence (black) and presence of 1 mM fragments. Left: chemical shift perturbations characteristic of fast exchange (FS-1092-06-B02). Middle: signal broadening and disappearance indicating intermediate exchange (FS-1255-03-C01. Right: signal intensity enhancement (11T-0672-02-B04), reflecting reduced solvent exchange upon binding. (B) Heatmap showing chemical shift perturbations and signal broadening induced by fragment binding. Rows represent hotspot residues (H80, C90, K91, W123); columns represent individual fragment hits. Coloured cells indicate significant CSPs (exceeding mean + 1σ threshold) (cyan) and signal broadening (pink); white cells indicate CSPs below significance threshold. (C) Hotspot residues (H80, C90, K91 and W123) mapped onto the AlphaFold-predicted structure of nsp10, shown as cyan sticks. (D) Binding affinities (K_d_) of the eight fragments estimated by MST. Values are given as K_d_ ± confidence interval, calculated by MO.Affinity Analysis from the variance of the fitted parameter returned by the Levenberg-Marquardt algorithm, corresponding to a 68% confidence level.

Multiple fragments perturbed overlapping regions, and residues H80, C90, K91 and W123 emerged as recurrent binding hot spots across several fragments (Figure 8). H80 lies on a solvent-exposed surface loop, and its imidazole sidechain (pKa ∼6.0-6.5) is sensitive to changes in local environment, so the origin of chemical shift perturbations should be interpreted carefully. The fragments perturbing H80 span a range of ionisation states and include neutral species, making systematic change in pH unlikely, and FS-1255-03-C01 additionally induced line broadening at this residue, characteristic of intermediate exchange and not expected from a shift in protonation equilibrium alone. S61 was considered a lower-confidence hotspot due to its weak signal intensity.

These hotspots map to functionally important regions of nsp10: H80 is located at the nsp10-nsp16 binding interface, where it participates in hydrogen bonding with nsp16 (PDB ID 5YN5); C90 is one of the four coordinating residues of zinc finger 1 (C74/C77/H83/C90) (Joseph, Saikatendu et al. 2006); K91 lies in the β-sheet subdomain adjacent to the nsp16 binding surface (PDB: 5YN5); and W123 is situated near zinc finger 2 (C117/C120/C128/C130), where its aromatic sidechain contributes to a hydrophobic surface feature (Joseph, Saikatendu et al. 2006). The clustering of fragment binding events at the two zinc finger regions and their adjacent protein-protein interaction interfaces is consistent with chemically specific binding rather than nonspecific surface adhesion, suggesting that these regions represent tractable sites for small-molecule intervention targeting nsp10-mediated protein-protein interactions.

In addition to these hotspots, residue L45, which undergoes rapid solvent exchange as confirmed by CLEANEX-PM experiments, showed signal enhancement upon addition of fragments 2W-0359-02-B01, 11T-0672-02-B04 and KS-1346-04-E12, suggesting that fragment binding reduces solvent accessibility at this position. L45 forms hydrogen bonds with C39 on nsp14 (Lin, Chen et al. 2021) and Q87 on nsp16 (Sk, Jonniya et al. 2020) in the respective complex, and L45A mutation has been shown to substantially reduce both nsp14 ExoN and nsp16 2’-O-MTase activity (Bouvet, Lugari et al. 2014). The observation that fragment binding sites overlap with these functionally important interface residues highlights the potential for fragment-derived compounds to interfere with nsp10-nsp14/nsp16 complex formation, thereby with viral replication.

Fragment FS-1255-03-C01 showed unusual binding kinetics, inducing signal broadening on several residues (T20, D64, S87 and R105) (Figure 8B), characteristic intermediate exchange kinetics despite a weak binding affinity by MST (K_d_ > 10 mM). For a fragment with millimolar range K_d_ and diffusion-limited association (k_on_ ≈ 10^7^-10^8^ M^-1^s^-1^), the dissociation rate constant k_off_ (= K_d_ × k_on_) would typically be on the order of 10^4^ s^-1^, placing the interaction in fast exchange. The observation of intermediate exchange instead implies that k_off_ is substantially slower than predicted, correspondingly k_on_ is anomalously slow, which may indicate a conformational gating mechanism that impedes dissociation.

Structural analysis of the 23 validated fragment hits revealed three predominant scaffolds: benzyl-azoles (eight hits), indole derivatives (six hits), and biphenyl compounds (four hits), together accounting for 78% of all hits (Figure 7D). Among the 20 validated hits, these three scaffold classes accounted for 80%. All three classes were represented among the eight fragments with quantifiable binding affinities, and no clear scaffold-dependent binding-site preference was observed, indicating that the nsp10 binding sites can accommodate various chemical structures. This scaffold diversity provides multiple starting points for further optimisation, with potential opportunities for structure-activity relationship studies through fragment growing, linking or merging approaches. Further structural characterisation of fragment binding modes would be required to guide rational optimisation strategies.

## Conclusions

This work provides the first solution-state NMR characterisation of unbound MERS-CoV nsp10. The folded core adopts the conserved coronavirus nsp10 fold and displays a predominantly rigid backbone, with localised flexibility in the loop connecting α3 and β3 and in the β-sheet of the lower subdomain. In contrast, no amide signals were detected for residues 10 to 22 under standard conditions. Signals for this region appear at reduced temperature and pH and give rise to CLEANEX-PM cross peaks, indicating rapid solvent exchange and high solvent exposure. Together with the absence of detectable ms conformational exchange for α2 residues adjacent to α1, these observations indicate that the N-terminal region, which forms the α1 helix in crystal structures of coronavirus nsp10, is disordered in unbound MERS-CoV nsp10 in solution. Fragment screening by ligand-observed ^19^F NMR identified 23 hits, of which 20 were confirmed by orthogonal methods, with chemical shift perturbations clustering near the zinc finger regions and their adjacent protein-protein interaction interfaces. The fragment hits reported here provide chemical starting points for optimisation towards compounds that interfere with nsp10-mediated complex formation.

Several aspects remain unresolved. The backbone assignments obtained at 283 K are tentative, since only limited sequential connectivity was observable at this temperature, although the conclusion of high solvent exposure does not depend on the specific assignment adopted. The structural basis of the difference between MERS-CoV nsp10 and SARS-CoV-2 nsp10, for which α1 is assigned and helical in solution, is not established by the present data. A conformational selection mechanism cannot be excluded, as a folded α1 conformation present at low population, or exchanging outside the timescale probed here, would not be detected. The fragments identified bind weakly, and for the weakest of these the affinities are constrained by the upper limit of the titration range. The perturbed residues indicate ligandable surfaces rather than a discrete pocket.

## Acknowledgements

The authors gratefully acknowledge financial support from the Medical Research Council (MRC), grant number MR/X013995/1. This work was supported by the Francis Crick Institute through provision of access to the MRC Biomedical NMR Centre. The Francis Crick Institute receives its core funding from Cancer Research UK (CC1078), the UK Medical Research Council (CC1078), and the Wellcome Trust (CC1078). We acknowledge use of the UCL School of Pharmacy Nuclear Magnetic Resonance Core Facility (RRID:SCR_027123). We thank Key Organics (UK) for a gift of the BioNET fluorine fragment library. We thank Dr Jake Williams for technical advice on the use of SPANR, and Dr Gary Thompson for developing a modified SPANR implementation that accepts NEF-format input in collaboration with Dr Williams. This work contains part of the doctoral thesis of Danni Dong.

## Notes

### Competing Interest Statement

The authors have declared no competing interest.

