## Supporting Information for "Solution structure, dynamics and fragment binding of unbound MERS-CoV nsp10"

### Supplementary sequences

*Before Ulp1 cleavage*

MGSSHHHHHHHHSSGLVPRGSHMASMSDSEVNQEAKPEVKPEVKPETHINLKVSDGSSEI
FFKIKKTTPLRRLMEAFAKRQGKEMDSLRFLYDGIRIQADQTPEDLDMEDNDIIEAHREQIGG
TMGNSSVLSLVNFTVDPQKAYLDFVNAGGAPLTNCVKMLTPKTGTGIAISVKPESTADQETY
GGASVCLYCRAHIEHPDVSGVCKYKGKFVQIPAQCVRDPVGFCLSNTPCNVCQYWIGYGC
NCD*

*After Ulp1 cleavage*

TMGNSSVLSLVNFTVDPQKAYLDFVNAGGAPLTNCVKMLTPKTGTGIAISVKPESTADQETY
GGASVCLYCRAHIEHPDVSGVCKYKGKFVQIPAQCVRDPVGFCLSNTPCNVCQYWIGYGC
NCD*

**Table S1. Main characteristics used in NMR data acquisition for nsp10 assignment and structure characterisation.**

| Experiments | Temperature (K) | Field strength (MHz) | Time domain data size (points) | | | Spectral width (ppm) | | | Number of scans | Recycle delay (s) | Mixing time (ms) |
|---|---|---|---|---|---|---|---|---|---|---|---|
| | | | F1 | F2 | F3 | F1 | F2 | F3 | | | |
| $^{1}$H, $^{15}$N SOFAST-HMQC | 298 | 600 | 160 | 2048 | - | 28 ($^{15}$N) | 16.0149 ($^{1}$H) | - | 8 | 0.1 | - |
| HNCACB | | 600 | 128 | 64 | 2048 | 80 ($^{13}$C) | 28 ($^{15}$N) | 16 ($^{1}$H) | 8 | 1.0 | - |
| HNCACO | | 600 | 64 | 64 | 2048 | 14 ($^{13}$C) | 28 ($^{15}$N) | 13.652 ($^{1}$H) | 16 | 1.0 | - |
| HNCO | | 600 | 64 | 64 | 2048 | 14 ($^{13}$C) | 28 ($^{15}$N) | 16 ($^{1}$H) | 8 | 0.2 | - |
| HNCOCACB | | 600 | 128 | 48 | 2048 | 80 ($^{13}$C) | 28 ($^{15}$N) | 16 ($^{1}$H) | 8 | 1.0 | - |
| $^{1}$H, $^{13}$C HSQC | | 800 | 256 | 3072 | - | 67 ($^{13}$C) | 16 ($^{1}$H) | - | 16 | 1.0 | - |
| H(C)CH-TOCSY | | 800 | 128 | 128 | 3072 | 7 ($^{1}$H) | 67 ($^{13}$C) | 16 ($^{1}$H) | 8 | 1.0 | 16.3 |
| (H)CCH-TOCSY | | 800 | 64 | 128 | 3072 | 67 ($^{13}$C) | 67 ($^{13}$C) | 16 ($^{1}$H) | 8 | 0.9 | 16.3 |
| $^{13}$C NOESY | | 700 | 256 | 64 | 3072 | 8.3 ($^{1}$H) | 61 ($^{13}$C) | 20 ($^{1}$H) | 8 | 1.0 | 160 |
| $^{15}$N NOESY | | 600 | 200 | 80 | 2048 | 12 ($^{1}$H ) | 29 ($^{15}$N) | 16 ($^{1}$H) | 4 | 1.0 | 80 |
| $^{1}$H, $^{15}$N SOFAST-HMQC | 283 | 800 | 192 | 3072 | - | 28 ($^{15}$N) | 18 ($^{1}$H) | - | 4 | 0.1 | - |
| BEST-HNCA | | 800 | 64 | 64 | 3072 | 24 ($^{13}$C) | 28 ($^{15}$N) | 18 ($^{1}$H) | 32 | 0.1 | - |
| BEST-HNCO | | 800 | 48 | 64 | 3072 | 14 ($^{13}$C) | 28 ($^{15}$N) | 18 ($^{1}$H) | 8 | 0.1 | - |
| BEST-HNCOCA | | 800 | 64 | 64 | 3072 | 24 ($^{13}$C) | 28 ($^{15}$N) | 18 ($^{1}$H) | 40 | 0.1 | - |
| BEST-HNCOCACB | | 800 | 112 | 60 | 3072 | 64 ($^{13}$C) | 28 ($^{15}$N) | 18 ($^{1}$H) | 32 | 0.1 | - |

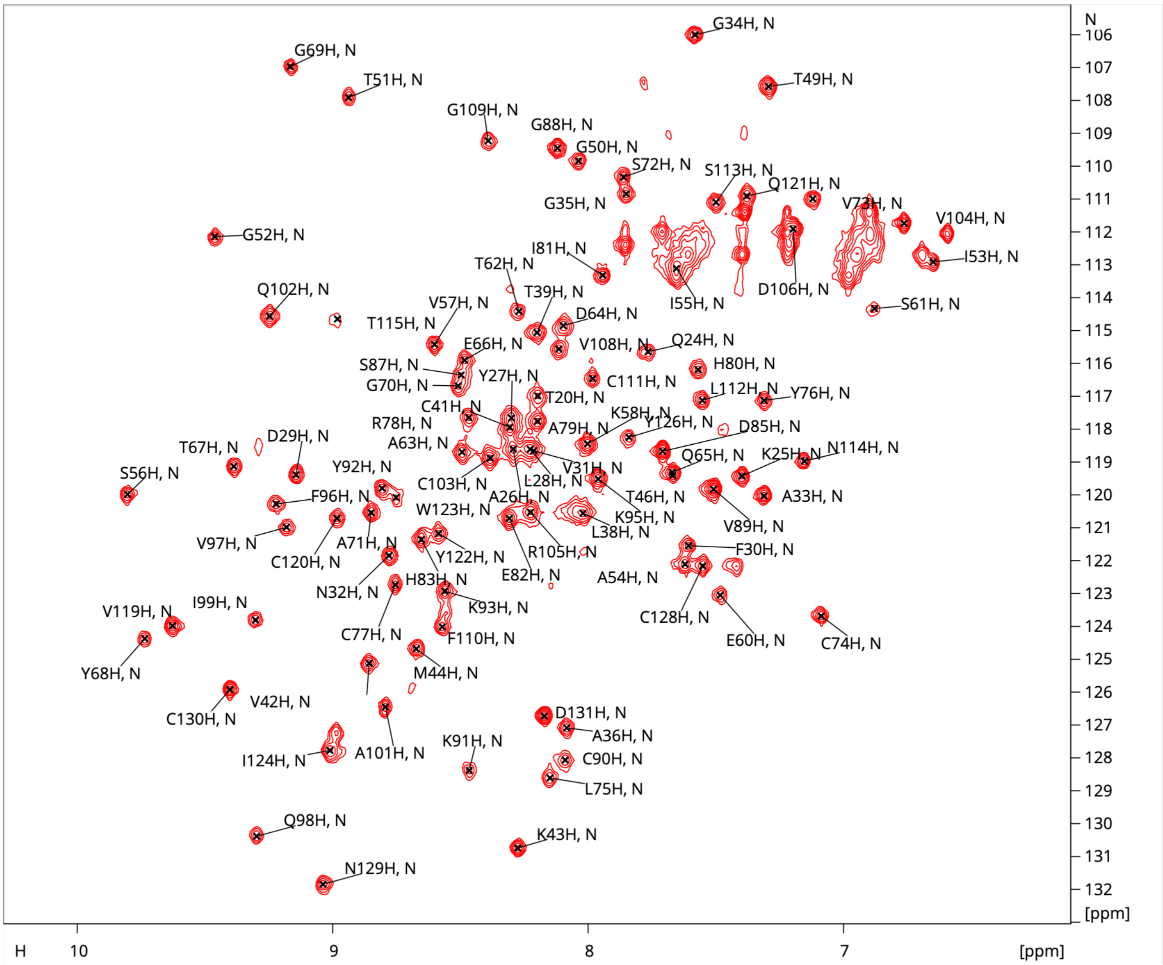


**Figure S1. $^1H$-$^{15}N$ SOFAST-HMQC spectrum with backbone amide assignments labelled.** Spectrum was recorded at 298 K, 600 MHz.

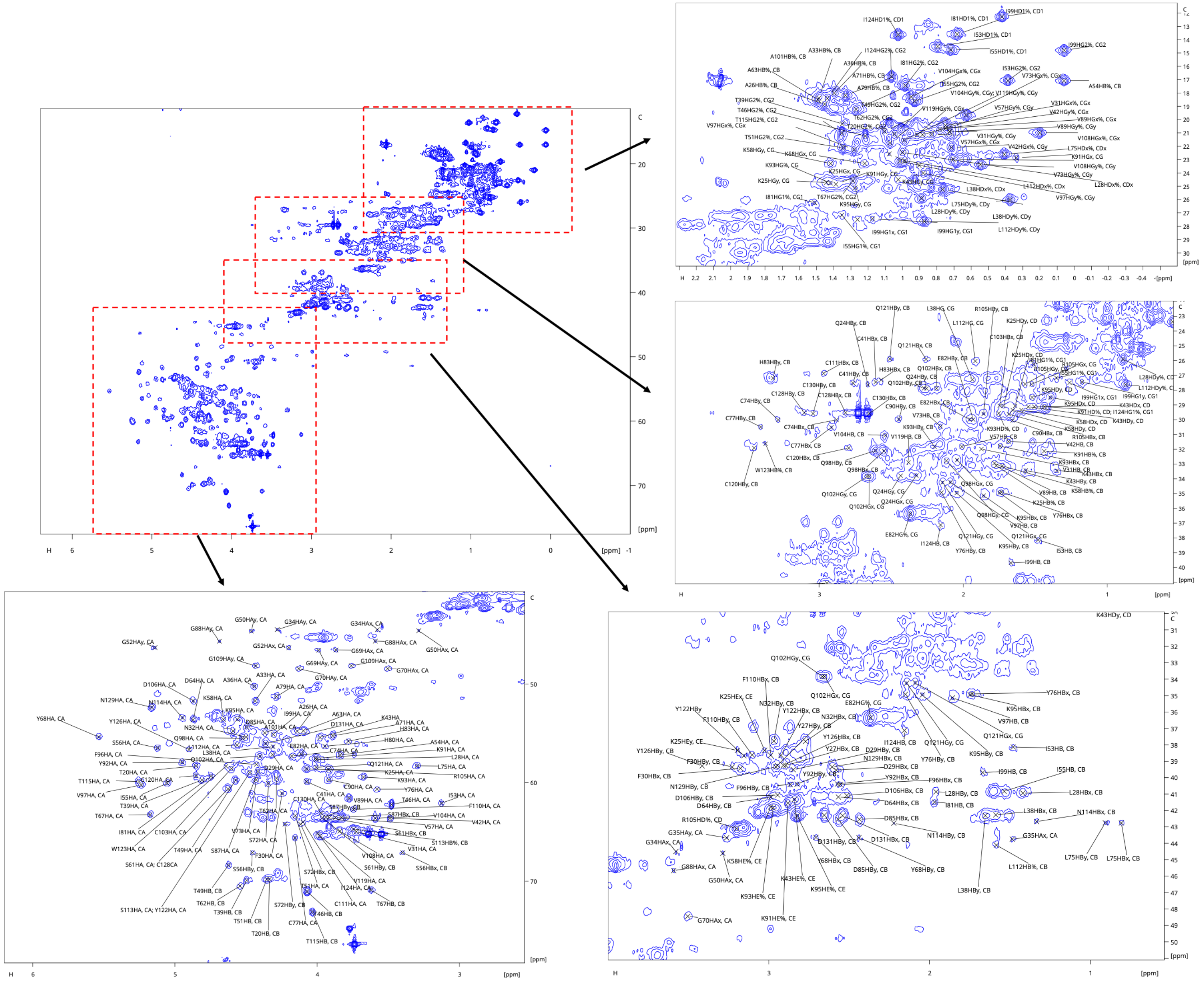

**Figure S2. ¹H-¹³C HSQC spectrum with sidechain assignments labelled.** Spectrum was recorded at 298 K, 800 MHz

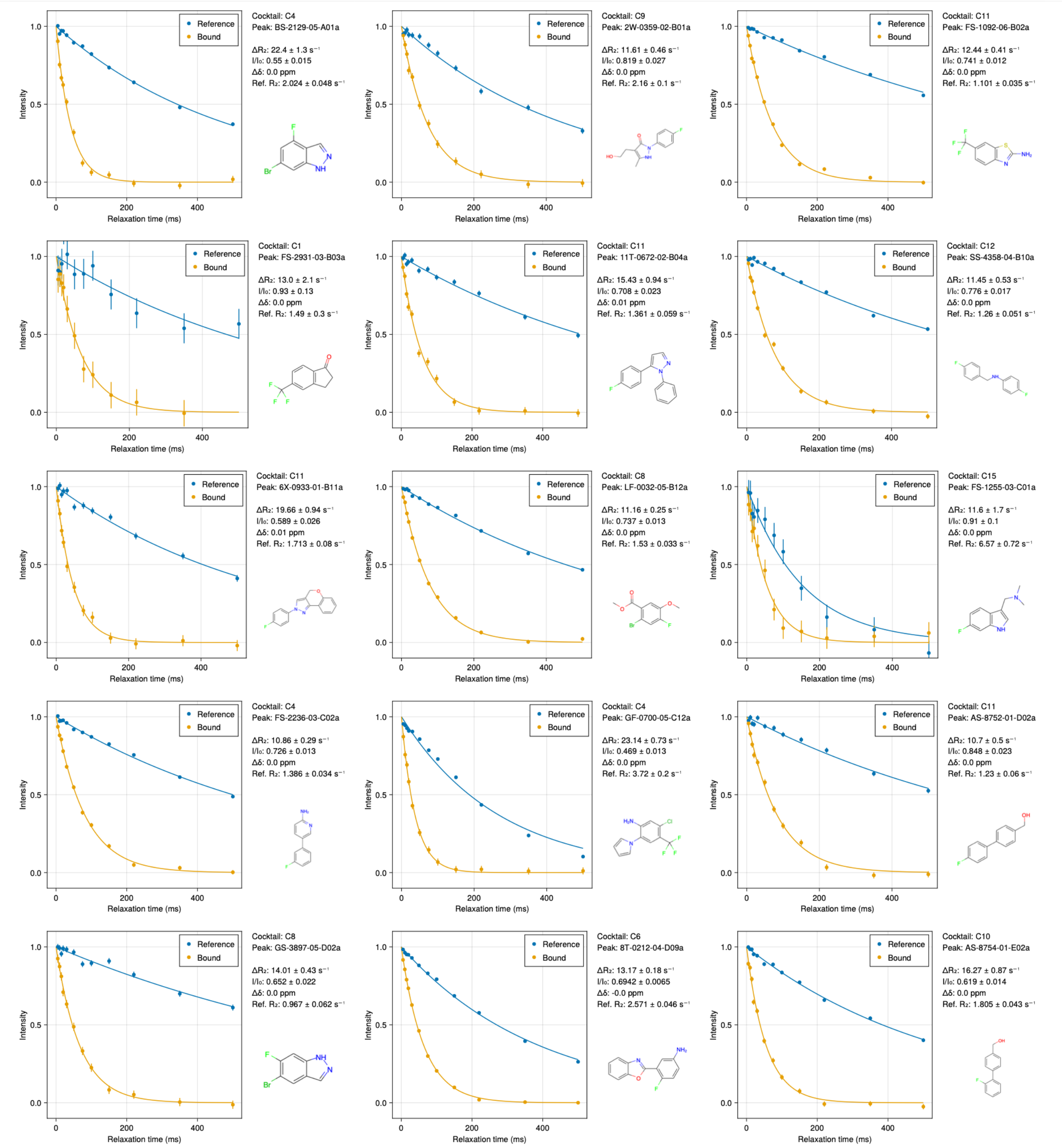


**Figure S3** (continues on next page)

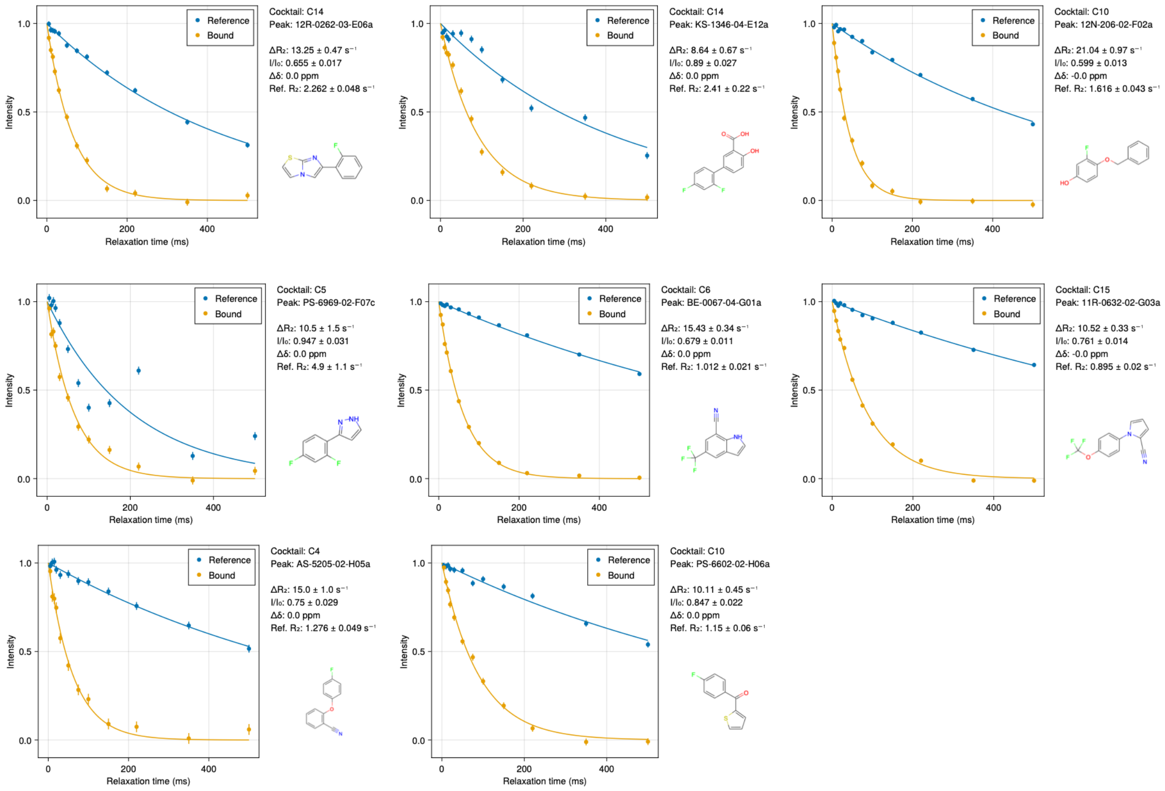


**Figure S3. $^{19}F$ $T_2$ relaxation decays for fragment hits (1 of 2) (600 MHz, 298 K).** Transverse relaxation decay curves measured in the absence (blue) and presence (orange) of nsp10, with relaxation delays from 5 to 500 ms. Solid lines are fits to a single-exponential decay. The fragment code and corresponding $\Delta R_2$ value are given above each panel.
